# Intrinsic plasticity underlies malleability of neural network heterogeneity

**DOI:** 10.1101/2025.06.09.658695

**Authors:** Daniel Trotter, Taufik Valiante, Jeremie Lefebvre

**Affiliations:** Department of Physics, University of Ottawa, Ottawa, Ontario, Canada; Krembil Research Institution, University Health Network, Toronto, Ontario, Canada; Division of Neurosurgery, Department of Surgery, University of Toronto, Toronto, Ontario, Canada; Division of Clinical and Computational Neuroscience, Krembil Brain Institute, University Health Network, Toronto, ON, Canada; Institute of Medical Science, University of Toronto, Toronto, ON, Canada; Institute of Biomedical Engineering, University of Toronto, Toronto, ON, Canada; Department of Electrical and Computer Engineering, University of Toronto, Toronto, ON, Canada; Center for Advancing Neurotechnological Innovation to Application (CRANIA), Toronto, ON, Canada; Max Planck-University of Toronto Center for Neural Science and Technology, University of Toronto, Toronto, ON, Canada; Department of Biology, University of Ottawa, Ottawa, Ontario, Canada; Department of Mathematics, University of Toronto, Toronto, Ontario, Canada

## Abstract

Diversity exists throughout biology, playing an important role in maintaining robustness and stability. The same is true of the brain, as has become increasingly apparent in recent years with the accumulation of datasets of unparalleled resolution. These datasets show widespread neural heterogeneity, spanning cells, circuits and system dynamics, marking it as an unavoidable component of the brain’s composition. Recent experiments found declines in heterogeneity amongst neurons may accompany pathological states. While heterogeneity has been linked to stability, robustness and increased computational potential, the loss of biophysical diversity was found conducive to the onset of seizure-like activity, suggesting an important functional role. Despite this, how changes in heterogeneity arise remains unknown. Oftentimes considered a static metaparameter resulting from solely genetic disposition, heterogeneity is, in fact, a highly dynamic property of biological networks arising from various sources. Here, we consider this through the lens of intrinsic plasticity, the activity-dependent modulation of neuron biophysical properties, which we propose allows the degree of biophysical diversity to fluctuate in time. Using a network of Poisson neurons endowed with intrinsic plasticity, we combine analytical and numerical approaches to measure the effect of input statistics on the excitability of individual cells, and how this translates into changes in network heterogeneity at the population scale. Our results indicate that, through intrinsic plasticity, diversity in synaptic inputs promotes heterogeneity in cell-to-cell excitability due to changes in the statistics of presynaptic firing rates, and network topology. In contrast, whenever the statistics of synaptic input between cells were too similar, intrinsic plasticity promoted the decline in heterogeneity. Further, we show that changes in heterogeneity can coexist with degeneracy in the firing rate between neurons. Taken together, understanding how input statistics affect neuronal network heterogeneity may provide key insights into brain function, resilience and the manipulability of neural diversity through intrinsic plasticity.

## I. INTRODUCTION

A common feature of many complex systems’ constitution is the presence of disorder or heterogeneity [1–8]. This is especially true for the brain, where the sources of heterogeneity are widespread [9, 10], with important consequences on neural circuits dynamics and function[Eq. (11)– Eq. (14)]. One of the most salient, and studied, forms of heterogeneity arises through diversity in neuronal excitability [15–17], manifested through neuron-specific, cell-to-cell variations in both the minimal intensity of a stimulus required to elicit a response (i.e., threshold), as well as in the magnitude of this response (i.e., gain) [18]. A confluence of both experimental and theoretical studies have demonstrated that excitability heterogeneity plays multiple important functional roles, such as optimizing coding capacity, promoting efficient learning and information flow, as well as reinforcing the resilience of neural circuits by stabilizing their dynamics away from pathological states [13, 17–22].

In all likelihood, because neuronal excitability is not static, excitability heterogeneity is also not static and modulated by *intrinsic plasticity* mechanisms that play an important role in learning and memory [23, 24]. Intrinsic plasticity is the post-synaptic counterpart to synaptic plasticity, whereby a neuron’s excitability (e.g. gain) is modulated by its past activity profile [Eq. (23), Eq. (25)– Eq. (28)], including background fluctuations and/or persistent changes in the input statistics (hereafter referred to as the “milieu”). Such modulation of excitability in an activity-dependent manner was first shown by blocking voltage-gated sodium channels to silence neocortical neurons, which triggered an increase in excitability via the upregulation of voltage-gated sodium channels and downregulation of potassium channels [23, 26, 29]. Experimentally, this malleability of excitability not only plays a key role in memory and learning [23, 30–33], but also in several pathological states of the nervous system [34] including epilepsy [23, 34], tinnitus [35–37], neuropathic pain [37, 38], and a whole host of neurodegenerative and neuropsychiatric conditions [39–46].

At a network level, the changes in excitability in pathological states like epilepsy can be seen to alter the diversity of excitability across the population. Indeed, our recent work shows that a loss of excitability diversity accompanies human [13] and rodent [47] epilepsy, and results in network instability that predisposes the brain to slipping into a seizure-like state [13, 48, 49]. However, how such a decline in heterogeneity arises remains unknown, and even of wider important is that how is the excitability diversity we see in the brain [Eq. (11), Eq. (12), Eq. (50)– Eq. (52)] maintained for the “healthy” function of the nervous system. Therefore, we pursue the hypothesis that intrinsic plasticity modulates neural circuit heterogeneity, through the regulation of the excitability of individual neurons. We consider this mechanism of particular candidacy as it aligns with recent research highlighting the importance of degeneracy in explaining the multiple pathways through which epilepsy occurs [53]. Indeed, if intrinsic plasticity enhances variability between neurons, either by diversifying their excitability profiles between each other or by accentuating pre-existing differences, the network’s heterogeneity will increase. In contrast, if intrinsic plasticity instead drives a convergence of excitability profiles between cells, this will manifest as a decline in excitability heterogeneity. Given the close relationship between diversity and stability [13, 48, 49], the ability of intrinsic plasticity to promote (or hinder) excitability heterogeneity might provoke important changes in dynamic stability, which may be both physiologically beneficial, or in the extreme predispose neural circuits to pathological states, such as those seen in epilepsy [13, 48, 49].

Inspired by our recent experimental work [47], we here pursue the novel hypothesis that intrinsic plasticity is a critical modulator of neural circuit excitability heterogeneity. This novel perspective is in stark contrast with the pre-existing notions that excitability heterogeneity is a static meta-parameter resulting mainly from genetic predisposition [Eq. (12), Eq. (14), Eq. (17), Eq. (51), Eq. (52), Eq. (54)–Eq. (56)]. Herein we propose intrinsic plasticity as a candidate mechanism by which intrinsic plasticity supports time-dependent changes in the heterogeneity expressed by neural populations, thereby influencing the stability of neural circuits. To explore these hypotheses, we use a widely used recurrent neural network of excitatory Poisson neurons with sparse, random connectivity, endowed with intrinsic plasticity. We characterize the dynamics of this model under varying stimulation conditions (e.g. step current, spike trains) and network architecture (e.g. topology and weights). We found that heterogeneity arises naturally among neurons from changes in input statistics, resulting from cell-to-cell variations in connectivity profiles. Most importantly, our result indicates that stimuli can be used to manipulate the level of heterogeneity in the network. The increase in heterogeneity was found to enhance network stability. In contrast, diversity decline was alternatively found to promote multistability. Importantly, we found that heterogeneity can coexist with degeneracy: networks of neurons exhibiting similar firing rates can still demonstrate excitability heterogeneity. Taken together, these results imply that heterogeneity is a malleable property of neural networks, supported by the topology of the network and imperative for maintaining stable neural network operation.

## II. METHODS

### A. Network model

We consider an excitatory network of Poisson point neurons with random connectivity. The neurons have a membrane potential, *u*_*i*_, where *i* = 1, …, *N* is the neuron index. The dynamics of the membrane potential for neurons in this network obey the following system of evolution equations,

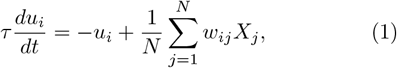

where 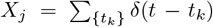 are Poisson spike trains, *τ* is the membrane time constant, and *w*_*ij*_ are synaptic weights. Spiking activity follows a non-homogeneous Poisson process, that is *X*_*j*_ → Poisson(*r*_*j*_), where *r*_*j*_ corresponds to the instantaneous firing rate of neuron *j*. This firing rate depends directly on the membrane potential through the excitability curve of that same neuron, that is *r*_*j*_ = *f* (*u*_*j*_) where *f* is defined by the sigmoid [48, 49],

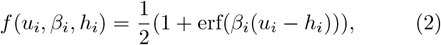

where *i* is the neuron index, and {*β*_*i*_, *h*_*i*_} are the slope and midpoint (or rheobase), respectively. A Poisson spike occurs if *p*_*spike*_ = 1 −exp(− *f* (*u*_*i*_, *β*_*i*_, *h*_*i*_)) is greater than a randomly generated uniform probability. The excitability curve hence sets the firing rate, *r*_*i*_, of neuron *i*.

Synaptic connectivity is assumed to be random with connection probability *ρ* = 0.15. For simplicity, synaptic weights are assumed to be equal across all realized connections i.e., *w*_*ij*_ = *w*_*o*_*/ρ >* 0 if a connection exists between neurons *i* and *j* and 0 otherwise. This choice is made for convenience, so that *E*_*N*×*N*_ [*w*_*ij*_] = *w*_*o*_. The network topology is constructed to accommodate any in- and out-degree distributions. To do so, we follow the steps described [57]. In summary, a vector of in-degrees, 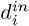, and out-degrees, 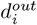, are drawn from the uniform distribution where *d*_*i*_ ≥ 0 ∀ *i* are the number of synaptic connections in/out of neuron *i*, where 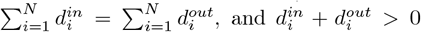. The connections are made by choosing a random neuron in the network, and matching all of its 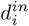 to available 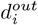. The available outward connections are placed in descending order, and the inward connections are then matched sequentially from left to right.

### B. Regulation of excitability via intrinsic plasticity

Thus far, the excitability described for each neuron in the network is static. To introduce intrinsic plasticity in this framework, we allow the parameters *β*_*i*_ and *h*_*i*_ in Eq. (2) to fluctuate in time to normalize the firing rate of individual neurons in an activity-dependent manner. Specifically, we allow the excitability curve of each neuron to adapt so that, over long time scales, its firing rate is drawn towards a value *r*^*o*^, hereafter referred to as the *homeostatic target rate*. For simplicity, this target rate is chosen to be equal for each neuron in the network. To implement intrinsic plasticity, Eq. (1) and (2) are complemented by an additional set of differential equations accounting for the dynamics of the parameters *β*_*i*_ and *h*_*i*_,

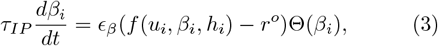

and

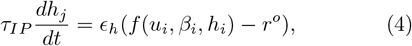

respectively. Combined, Eqs. (1), (3) and (4) form a system of 3*N* differential equations, in which the parameters defining the excitability of individual neurons adapts to fluctuations in its membrane potential. The constant *τ*_*IP*_ refers to the time constant of intrinsic plasticity, and is assumed to be slow compared to the dynamics of the membrane potential in Eq. (2) i.e. *τ*_*IP*_ ≪ *τ*. The parameters *ϵ*_*β*_ and *ϵ*_*h*_ correspond to learning rates (*ϵ*_*β*_ *<* 0 and *ϵ*_*h*_ *>* 0), scaling the direction of the rate of change, while compensating for the vastly different range of values the parameters *β*_*i*_ and *h*_*i*_ can take. The function Θ in Eq. (3) is defined by Θ(*β*_*i*_) = 1 whenever 0 *< β*_*i*_ *< β*_*max*_ and zero otherwise, and is included to ensure that the slopes *β*_*i*_ remain positive, and bounded within a physiologically plausible range, where *β*_*max*_ = 50.

### C. Excitability and degeneracy

To characterize how intrinsic plasticity influences the activity of individual cells in Eq. (1), one can examine the asymptotic dynamics of the parameters *β*_*i*_ and *h*_*i*_ for an arbitrary neuron *i*. Since the membrane potential *u*_*i*_ is a stochastic state variable (due to Poisson spiking), it is more appropriate to consider the temporal averages of Eq. (3) and (4), that is

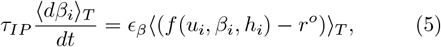

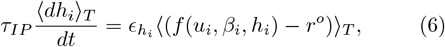

where ⟨·⟩_*T*_ represents an average over a long time window i.e., *T* ≫ *τ*_*IP*_ and *T* ≫ *τ* and where we assumed Θ(*β*_*i*_) = 1. At equilibrium, these equations possess steady states 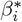 and 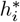 that satisfy the same implicit expression

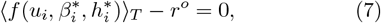

which simply indicates that asymptotically, the average individual firing rate of a neuron *i* will converge towards the homeostatic target rate *r*_*o*_. This can can equivalently be written for *T* → ∞ as,

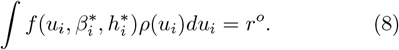

Note that one obtains an identical expression for each neuron in the network. The above expression depends directly on the a priori unknown probability density function *ρ*(*u*_*i*_) for the membrane potential of neuron *i*, as defined in Eq. (1). Determining this distribution exactly represents a challenge, but approximations are possible. Indeed, in the limit of high firing rates, and assuming that spiking activity in the network is uncorrelated, one may replace the spike trains, *X*_*j*_, in Eq (1) by 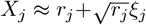, where *r*_*j*_ is the firing rate of neuron *j*, and *ξ*_*j*_ is a Gaussian white noise process such that ⟨*ξ*_*j*_(*t*)*ξ*_*k*_(*s*) ⟩ _*T*_ = *δ*_*jk*_(*t* −*s*) [58]. Using these approximations, Eq. (1) becomes,

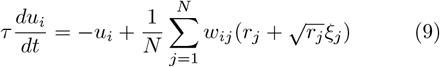

As per the steady state analysis of the intrinsic plasticity equations performed above, we know that asymptotically the individual firing rates *r*_*j*_ will converge towards their homeostatic target rate *r*^*o*^ (i.e., Eq. (7)). Hence,

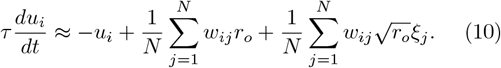

One may simplify this expression further in the case where the degree distribution is relatively uniform, and recalling synaptic weights are assumed to be the same for all connections i.e., *w*_*i,j*_ = *w*_*o*_ with probability *ρ*. Note that we will break this assumption later on. In this case, in the limit of large *N*, one may write

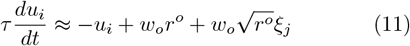

The above approximate equation has been fully decoupled from other neurons in the network, and further indicates that the dynamics of the membrane potential are identical for all neurons *i*. Dropping the index *i* for clarity, the membrane potential dynamics in Eq. (11) correspond to a simple Ornstein-Uhlenbeck process whose probability density function *ρ*(*u*) is normally distributed for with mean

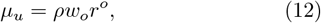

and variance

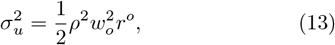

where we assume *τ* = 1. With an approximation for the probability density function 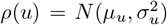 in hand, we can now return to Eq. (8) and compute the integral over the excitability curve on the right hand side

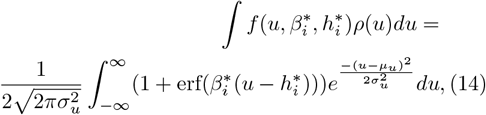

which can be separated into two integral terms,

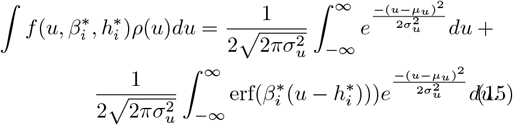

The first term of this equation resolves to 1*/*2. The second term can be solved using the known integral solution of a normal distribution with an error function [59], yielding

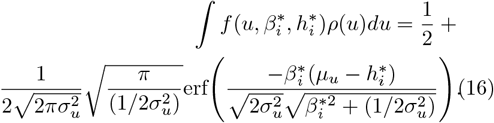

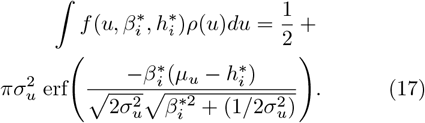

Simplifying this equation, we obtain an expression for the average firing rate of individual neurons, which at equilibrium must correspond to the homeostatic rate as per Eq. (7), that is,

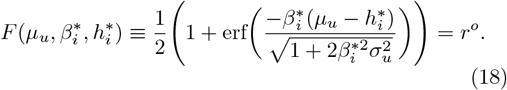

Equation 18 defines a self consistent, yet implicit expression for values of 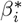 and 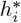 reached at equilibrium, which result from Eqs. (3) and (4). It further confirms that the shape of the excitability curve (which is parametrized by the parameters 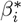 and 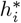) depends directly on the statistics of the membrane potential, and thus that the excitability of individual neurons changes in an activitydependent manner to normalize the neuron(s) firing rate. As defined, the corresponding equilibrium equations (i.e., which are identical; see Eq. 7) form an underdetermined system and, by extension, the model displays *degeneracy*. That is, Eq. (18) admits multiple equivalent solutions, where multiple combinations of 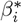 and 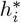 satisfy the equilibrium conditions of the intrinsic plasticity equations. That is, within the available *h*−*β* parameter space there exists a set of equivalent pairs of 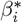 and 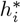 that parameterize an excitability curve with the same qualitative excitability profile. These define a curve, which we call the *degeneracy curve h*^∗^(*β*^∗^), towards which values of *β* and *h* will converge to maintain the neuron at a set homeostatic target firing rate (see Fig (1C)).

**FIG. 1.**
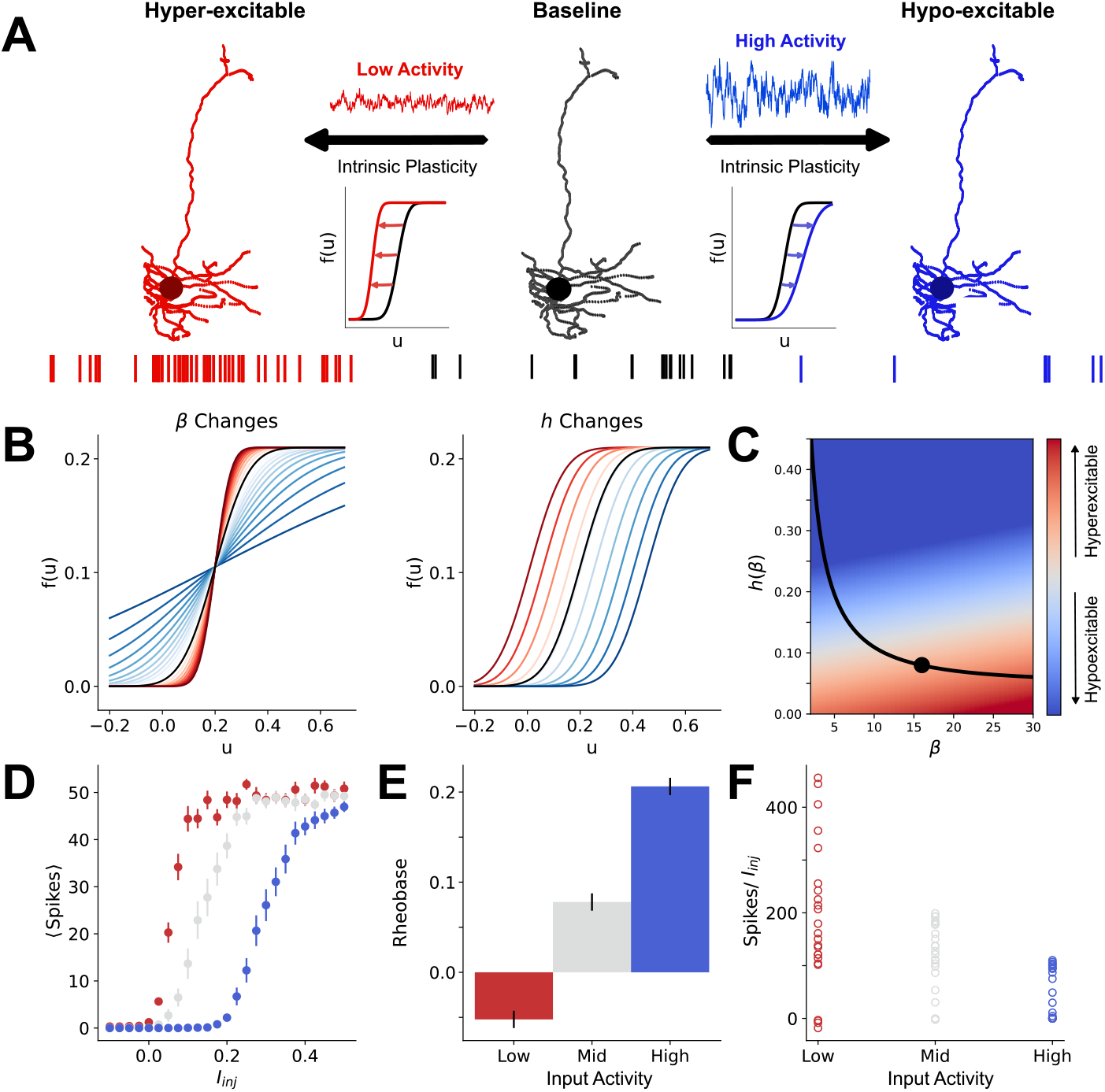
Intrinsic plasticity regulates neuron excitability in response to input statistics. **A)** For a given neuron, its excitability is altered by the input statistics it receives via intrinsic plasticity. Changes in input statistics can result from multiple factors, including connectivity and weights. In response to an increase in activity (such as from a high amount of input; blue), a neuron becomes hypoexcitable. Conversely, a low activity (red) will yield hyperexcitability. Both of these directions of change are facilitated by intrinsic plasticity. **B)** Intrinsic plasticity adapts the slope, *β* (left) and rheobase, *h* (right) of the excitability in response to high (blue) and low (red) input activity. **C)** The slope and rheobase the excitability adopts are directly influenced by the intensity of input received, where high (blue) intensity yields hypoexcitable low slopes and high rheobase values, and low (red) intensity the opposite. The black line is the degeneracy curve, and the black circle acts as an exemplar initial parameter set for a given neuron. When the input intensity changes to lower or high activity, the neuron will adopt new parameters via intrinsic plasticity that are more hyperexcitable (low activity) or hypoexcitable (high activity), the relative parameter spaces for which are demonstratively shown as a heat map. In a simulated network of *N* = 100 excitatory neurons, the varied statistics of their presynaptic firing rates will cause a heterogeneous population of excitabilities to emerge. By assigning thresholds, these excitabilities can be grouped into three broad categories of the amount of activity the neurons receive (low (red), mid (grey), and high (blue)). Correspondingly, this results in neurons that are hyperexcitable (red) and hypoexcitable (blue) with the mid group having excitability between these extremes. **D)** This excitability is shown as the average number of spikes produced by neurons in each grouping when stimulated by step currents, *I*_*inj*_, for *t* = 700 ms intervals with increasing current amplitude (see Methods). Error bars are SEM. The spikes evoked by the step current are recorded and reflect the rheobase, *h*, **E)** and gain, *β*, **F)**.

The degeneracy curve arises directly from Eq (18), where 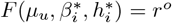. Assuming that 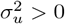 and performing the appropriate rearrangement to isolate *h*_*i*_, we hence obtain an expression for the degeneracy curve,

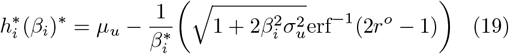

The degeneracy curve depends on the homeostatic target firing rate, as well as the mean and variance of the neuron’s membrane potential, which themselves depend on the weights and firing rates within the network (i.e., Eq. (12 and (13)). The presence of degeneracy in the plasticity of our model differentiates it from other adaptation models [60–66] as there are equivalent parametric pairs that can arise from inputs with the same statistics. Further, in cases where the neurons have non-identical input statistics (from e.g. degree distribution, connections weights) the presence of degeneracy allows for elevated levels of heterogeneity in the network. This degeneracy allows neurons to have different parameterizations, increasing the available parameter space beyond what a non-degenerate model can access. This is particularly important, as it is consistent with neuronal adaptations that occur via the complex interaction of multiple biophysical properties [67, 68].

### D. Mean field analysis

To formally relate intrinsic plasticity and heterogeneity at the network scale in Eq. (1) we leverage an approach inspired by mean field theory [48, 49]. Taking the mean of Eq (1) yields

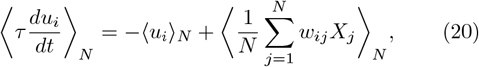

where ⟨·⟩ _*N*_ = E_*N*_ [·] corresponds to the ensemble average taken across all neurons in the network. Recalling that in a regime of high firing rates and uncorrelated spiking, 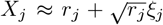 [58], and that *r*_*j*_ = *f* (*u*_*j*_, *β*_*j*_, *h*_*j*_), one may write

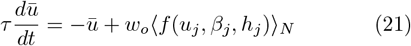

since ⟨*ξ*_*j*_⟩ _*N*_ = 0. In the above, we assumed independence between the firing rates and synaptic weights i.e. *E* _*N*_ [*w*_*ij*_*r*_*j*_] = *E* _*N*_ [*w*_*ij*_] *E* _*N*_ [*r*_*j*_], which holds whenever the degree distribution is relatively uniform, and introduced the population mean membrane potential (i.e., the mean field) defined by

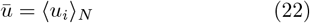

Equation (21) warrants further examination of the population-averaged response function, defined by

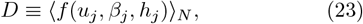

which for *N* → ∞ may be written as

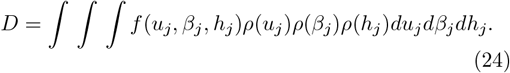

The above expression depends on the priori unknown membrane potential probability density function *ρ*(*u*_*i*_) (which may differ between neurons), as well as the distributions for the slopes *ρ*(*β*_*i*_) and rheobases *ρ*(*h*_*i*_) found across the network.

To the best of our knowledge, Eq. (24) does not possess a well-defined analytical solution for arbitrary distributions. Previous work has however shown that whenever *ρ*(*u*_*i*_) and *ρ*(*h*_*i*_) are normally distributed, Eq.(24) is also a sigmoid. The slope and rheobase of this resulting sigmoid depend on the mean and variance of the distributions *ρ*(*u*_*i*_) and *ρ*(*h*_*i*_), and hence scales with network heterogeneity in those parameters [48, 49]. Motivated by these findings as well as robust numerical observations (i.e., see Figs 2,5), we assume the population-averaged response function in Eq. (23) is also sigmoidally shaped and write

**FIG. 2.**
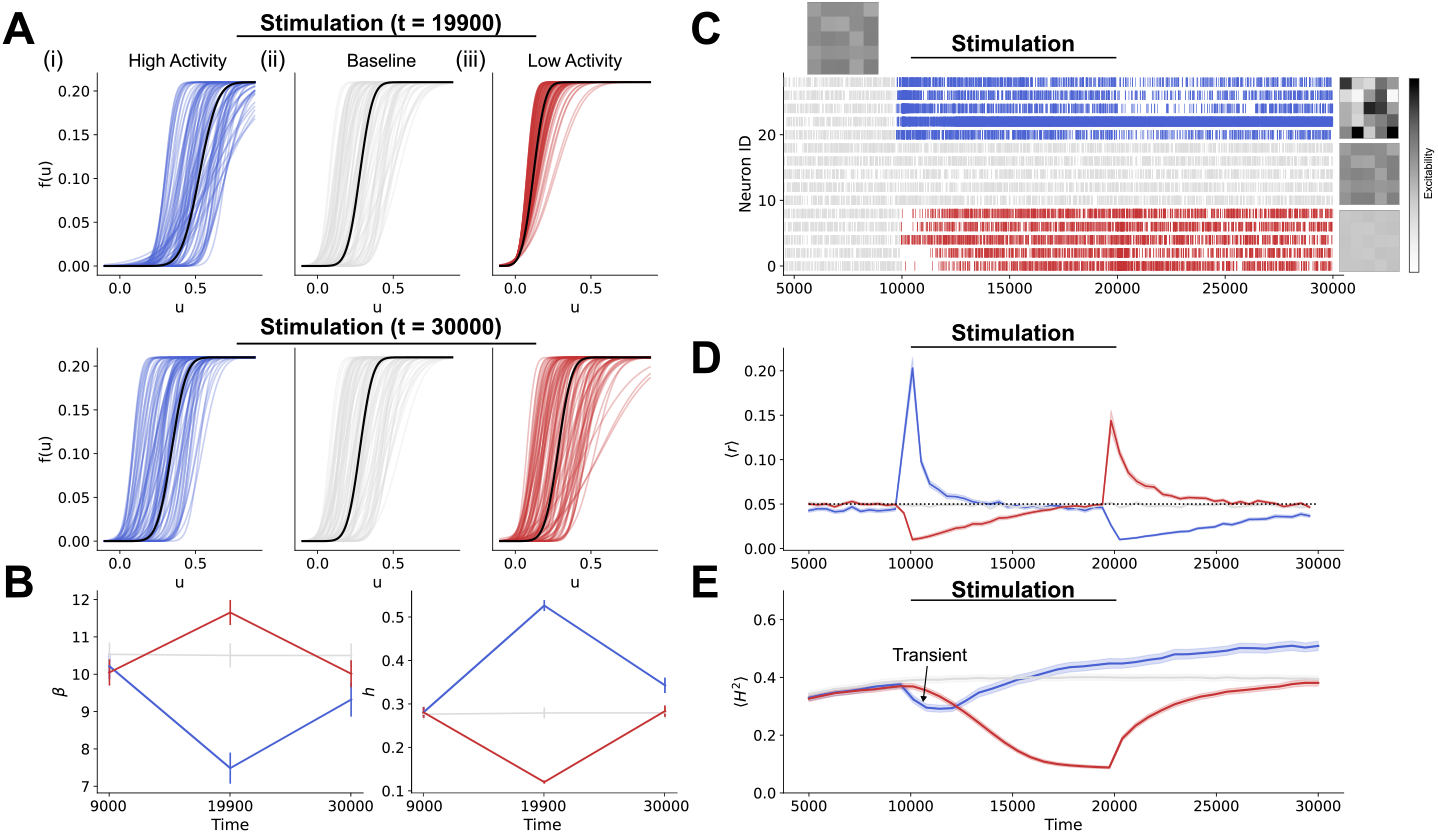
Persistent high and low activity states modulate both excitability and heterogeneity via intrinsic plasticity. **A)** Top: The coloured curves represent the excitabilities obtained by the *N* = 100 individual neurons connected with mean degree 15, and the black curve is the mean network response function, ⟨*f* (*u, β*_*eff*_, *h*_*eff*_) ⟩_*N*_, from simulations of a network initialized with random *β*_*o,i*_ and *h*_*o,i*_ at Top: *t* = 19900 ms following 9900 ms of exposure to stimulation of (i) persistent high activity, +*I*_*o*_ (blue), (ii) continued baseline activity, *I*_*o*_ = 0 (grey), and (iii) persistent low activity, − *I*_*o*_ (red). Bottom: plots of the excitabilities at *t* = 30000 ms, 10000 ms after exposure to persistent high (i) and low (iii) activity has stopped. **B)** The average population value of *β* (left) and *h* (right) before (*t* = 9000), during (*t* = 19900) and after (*t* = 30000) persistent activity. The error bars are SEM. **C)** A raster of the firing rate of a random sample of *n* = 5 neurons from the network stimulated with persistent high activity (blue), baseline activity (grey) and persistent low activity (red). The heatmaps are schematic representations of the heterogeneity in excitability obtained in the simulated networks. **D)** The population average firing rate, *r*, throughout the simulations. The black bar indicates when stimulation is applied to provide persistent high (blue) and low (red) activity. The horizontal dashed line is the target rate, *r*^*o*^ = 0.05. **E)** The heterogeneity level in the network, calculated as the population average Hellinger distance, ⟨*H*^2^⟩ _*N*_, (see Methods) is calculated as the pairwise average across all neurons. The black bar indicates exposure to stimulation. A window of transient behaviour, indicated by the arrow on the plot, occurs with the onset of high activity (blue) resulting in a temporary decline in heterogeneity as intrinsic plasticity is induced.

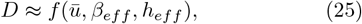

where *β*_*eff*_ and *h*_*eff*_ are the effective slope and rheobase of the population averaged response. The value of these parameters reflected the heterogeneity in excitability found in the network, and will be determined numerically. It is indeed known that increased heterogeneity will generally decrease the slope *β*_*eff*_ of the mean response function [48, 49], effectively making the system behave more linearly. Combining the steps above, we may recast Eq. (21) and obtain the dynamics of the mean field,

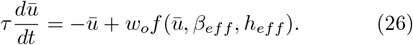

**Steady state.** At equilibrium, intrinsic plasticity and network dynamics have both stabilized. There, the parameters 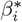 and 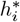 lie on the degeneracy curve (i.e., Eq. (19)) and one has *f* (*u*_*j*_, *β*_*j*_, *h*_*j*_) ≈ *r*^*o*^, ∀*j* as per Eq. (7). Hence, Eq. (23) reduces to

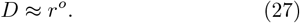

Thus, within the scope of the approximations above, the steady state of Eq. (26) is found to be

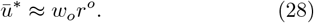

The steady state *ū*^∗^ above is furthermore unique. Indeed, since the derivative of Eq. (2) with respect to *u*_*i*_ is always positive, Eq. (7) possesses a single root. We further highlight that this steady state does not depend explicitly on the excitability of individual neurons in the network, and results directly from homeostasis, by which the firing rate of each cell is normalized.

**Stability.** To characterize the effect of excitability heterogeneity on network stability, we rely on an adiabatic approximation. We first assume that under the action of the intrinsic plasticity equations, which operate on a slower time scale, the system has reached equilibrium: neurons adopt their individual excitability profiles (i.e., 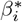 and 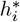), leading the network towards its steady state activity *ū*^∗^ defined above. Over short time scales however (≪ *τ*_*IP*_), the excitability profile of individual neurons can be assumed to be constant: the value of the parameters *β*_*i*_ and *h*_*i*_ do not change, even the membrane potential of neurons is perturbed. As such, in this adiabatic regime, the stability of the system depends only on the membrane potential (i.e., Eq. (1)), and we may thus restrict our analysis to the mean field equation. While approximate, this approach allows us to relate heterogeneity with stability resulting from the parameters *β*_*eff*_ and *h*_*eff*_ over short time scales.

Systems such as those in Eq. (26) are known to exhibit multistability. The following steady state equation

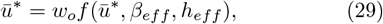

may indeed admit one or three solutions, depending on the values of the parameters *β*_*eff*_ and *h*_*eff*_ [Eq. (69)?, Eq. (70)]. Linear stability analysis indicates that the stability of these fixed points is determined by the following characteristic equation for *λ* ∈ ℝ,

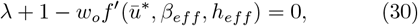

with 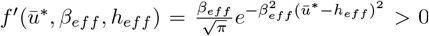. For the fixed point defined in Eq. (28), this equation becomes

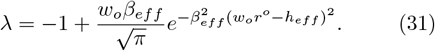

The above equation indicates that stability of the steady state *ū*^∗^ decreases whenever *β*_*eff*_ increases and *h*_*eff*_ approaches the value *w*_*o*_*r*^*o*^, conditions in which the second term on the right hand side of Eq. (31) becomes large. We note that this analysis is only valid over short time scales. Over longer time scales, perturbations of the membrane potential will be compensated for by intrinsic plasticity, driving the system back to its baseline steady state *ū*^∗^ defined above. Thus, strictly speaking, the steady state *ū*^∗^ is stable over long time scales, but allows for brief periods of instability.

### E. Table of parameters

### F. Quantifying heterogeneity

To discuss the heterogeneity of the excitability curves within the network, we first require a metric to quantify it. In choosing a metric for this, we capitalize on the fact that the derivative of the excitability curves is normally distributed, for which there exists a number of established similarity metrics [71–73]. The derivative of the excitability curve is given by,

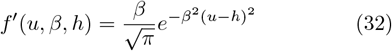

where we have dropped the index *i* for convenience, and further adopts the form of a normal distribution of mean, *µ*, and variance, *σ*^2^ which can be written in terms of *β* and *h*. Specifically, *µ* = *h* and *σ*^2^ = 1*/*(2*β*^2^). We use these values to calculate the Hellinger distance, *H*^2^ (Eq (36)). The Hellinger distance is a type of f-divergence used to define the difference between two probability distributions [74]. We chose it as our heterogeneity metric because, leveraging the normally distributed shape of the derivative of the excitability (Eq (32)), it can be fully resolved in terms of *β* and *h* (Eq (36)) allowing for a more direct relationship to our excitability. Moreover, the Hellinger distance is bounded between 0 and 1, and is thus a readily interpretable measure of the network heterogeneity where 0 corresponds to fully homogeneous and 1 would indicate heterogeneity with no cell-to-cell overlap. Between any two probability distributions *P* and *Q*, the Hellinger distance is defined on measure space *χ* as,

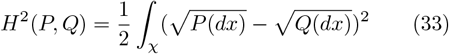

for a discrete distribution *P* = (*p*_1_, …, *p*_*k*_) and *Q* = (*q*_1_, …, *q*_*k*_), this is,

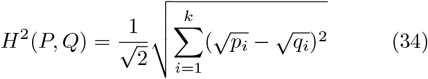

**TABLE 1.** Model parameters.

| Symbol | Definition | Value |
| --- | --- | --- |
| $u_i$ | Membrane potential of neuron $i$ | N/A |
| $\beta_i$ | Slope of the excitability curve | 0.1-50 |
| $h_i$ | Rheobase of neuron $i$ | 0-0.7 |
| $N$ | Number of neurons in the network | 100 |
| $r^o$ | The target firing rate of the neurons | 0.05 |
| $\epsilon_\beta$ | The learning rate of $\beta$ | -0.05 |
| $\epsilon_h$ | The learning rate of $h$ | 0.001 |
| $\tau$ | Membrane time constant | 1 |
| $r_i$ | Firing rate of neuron $i$ | N/A |
| $dt$ | Integration time step | 0.1 |
| $w_{ij}$ | Weight of connection between neuron $i$ and $j$ | 0-0.4 |
| $w_o$ | Fixed connection weight of all $w_{ij}$ unless otherwise stated | 0.21 |
| $\rho$ | Connection probability | 0.15 |
| $X_i$ | Poisson spike train | N/A |

This measure is bounded between 0 and 1, where the maximum distance 1 is achieved if there is no positive overlap whatsoever between the two distributions. For the special case where the distances are calculated between two normal distributions, 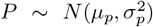 and 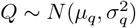, we can write *H*^2^ as,

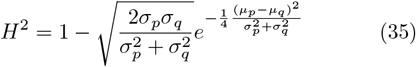

where *µ*’s and *σ*^2^’s are the mean and variance of the respective distributions, as noted earlier, these can be written in terms of *β* and *h* for the excitability curves. This then yields,

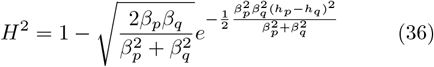

where *β*_*p*_, *β*_*q*_ *>* 0 ∀ *p, q*. To quantify the heterogeneity within the network, this value is calculated pairwise for each neuron, and the result is then averaged.

### G. Network simulations

#### 1. Manipulating network heterogeneity: stimulation

The input statistics of each neuron regulate the network’s level of heterogeneity by way of the membrane potential mean and variance (Eq (12) and (13)). As such, we would expect that heterogeneity will be modulated by exogenous stimulation. To explore how different types of stimulation affect network heterogeneity, we apply stimulation, *S*_*i*_, to every neuron in the network such that Eq (1) becomes,

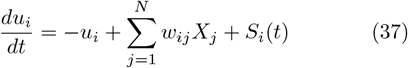

where *S*_*i*_ is representative of different kinds of stimulation:

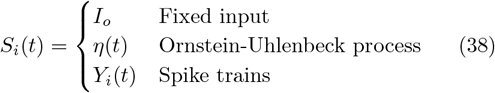

These correspond to different stimulus types regularly observed in experimental protocols and *in vivo* [75].

Starting with the case of fixed input, *S*_*i*_ corresponds to regimes of persistent high or low activity whereupon every neuron receives the same injected input. This kind of persistent stimulation is modelled in Section III B, where three different simulations are performed: one with persistently high activity where *I*_*o*_ *>* 0, one with persistently low activity where *I*_*o*_ *<* 0, and a baseline case where *I*_*o*_ = 0 ∀ *t*. The networks are initialized with random values of *β*_*i,o*_, *h*_*i,o*_ from uniform distributions of *β* and *h* for neurons *i* = 0, …, *N*.

At the end of these simulations, the obtained parameterizations of the excitability curves (e.g. *β*_*i,f*_, *h*_*i,f*_) for each of the three networks are kept to investigate the dynamics these curves would evoke in a static network. That is, *without* intrinsic plasticity to adjust the excitability curves, what dynamics may result from the new levels of excitability and homogeneity. This no-plasticity version of the network is then simulated with an Ornstein-Uhlenbeck process, *η*, such that every neuron receives the same slow-varying signal. This variety of stimulation allows us to investigate the influence of heterogeneity on the stability of the network dynamics.

Finally, in the case where the network is stimulated by impinging spike trains each train, *Y*_*i*_, *i* = 1, …, *N*, is independent of the *N* − 1 other trains and has a rate, *r*_*i,s*_, drawn from a uniform distribution of rates. The rates are non-identical between neurons with the objective of increasing the heterogeneity in the population. Simulations using this kind of stimulation are initialized with random *β*_*i,o*_, *h*_*i,o*_, the spike trains are applied over a window Δ*t*, to induce heterogeneity in the network.

#### 2. Manipulating network heterogeneity: In-network properties

In addition to exploring the effect of stimulation, we sought to investigate the effect of the network’s properties on excitability and heterogeneity. Specifically, we consider the impact of i) the topology and ii) the connectivity weights. These quantities are of particular interest due to their role in setting the mean (see Eq(12)) and variance (Eq(13)) of the membrane potential. To do this, rather than stimulating, we alter the input statistics of the neurons by changing i) the degree distribution of the topology or ii) the weights.

Specifically, to explore the effect of the topology on the network heterogeneity, we fix the mean connectivity to 15%, but replace the distribution the number of connections is drawn from (see Methods - Section (II A)) with a uniform one. In subsequent simulations, the variance of the uniform distribution is increased to observe the effect on network heterogeneity. For all of these simulations, the weights are maintained at a constant value, *w*_*i,j*_ = *w*_*o*_.

For the contribution of the weights, the topology is kept to the initial degree distribution with 15% random connectivity. However, the weights are no longer constant (*w*_*i,j*_ ≠ *w*_*o*_). Instead, the weights are drawn from uniform distributions and multiplied by the connections (*C*_*ij*_ = 0 or 1), to obtain the realized weight matrix *W*. As the network is all excitatory, the weights are all chosen to be *w*_*ij*_ ≥ 0.

In both cases, it is of interest to know what level of heterogeneity these in-network properties create and their ability to retain any induced heterogeneity. Thus the simulations for each case are first allowed to run until a baseline level of heterogeneity is achieved, after which spike trains of different rates (*Y*_*i*_ in Eq (37)) are applied for an interval *t* = 3000 to induce elevated heterogeneity in the network. When the stimulation ends, it can then be observed in each framework if, and to what extent, the induced heterogeneity is successfully retained.

## III. RESULTS

### A. Intrinsic plasticity and neural excitability

To explore the relationship between intrinsic plasticity and heterogeneity, we considered a simple and well-studied [60, 76] excitatory network of Poisson neurons with sparse, random connectivity exposed to multiple stimuli. We endowed the neurons within this network with a phenomenological model of intrinsic plasticity, whereby the excitability of individual neurons changes in an activity-dependent manner. Similar to other forms of adaptation [Eq. (60)–Eq. (64), Eq. (77)], homeostatic intrinsic plasticity was implemented by allowing the slope, *β*, and rheobase, *h*, of each neuron’s excitability curves to change over a slow timescale compared to neural activity (*τ* ≪ 1) (see Methods) as a function of membrane potential fluctuations (see Fig (1A,B)), while preserving a target homeostatic firing rate. The asymptotic values of *β* and *h* - which define the shape of the excitability curve - correspond to the steady states of the intrinsic plasticity equations (Eq. (19); see Methods).

A key difference between the myriad of other models of neural adaptation [Eq. (60)–Eq. (66), Eq. (76), Eq. (77)] and ours stems from the presence of degeneracy, a phenomenon where multiple unique sets of solutions can produce a highly similar performance. Importantly, this differs from redundancy, where multiple sets of the same elements perform the same function [53, 78, 79]. While long understood in theory, it is only more recently that degeneracy has been understood as a pervasive property of biological systems, including in the brain where neurons with widely varying ion channel conductances can nonetheless exhibit remarkably similar electrophysiological properties [Eq. (67), Eq. (68), Eq. (80)–Eq. (82)]. This is particularly true for excitability, as multiple forms of contextual, environmental, and neuromodulatory perturbations may cause changes in input statistics to trigger intrinsic plasticity, ranging from variations in synaptic dynamics, synchrony, input amplitude and position, shunting inhibition, total conductance, depolarization state, and fluctuations in the membrane potential [18]. Leveraging this in our model, we implement degeneracy by allowing the neurons to possess different excitability profiles while sharing a common target firing rate. This arose from the intrinsic plasticity equations, which form an underdetermined system, allowing for multiple equivalent solutions. By this, numerous combinations of *β* and *h* satisfy the equilibrium conditions of the intrinsic plasticity equations, which define a curve which we call the *degeneracy curve, h*(*β*). Over time *β* and *h* converge to the degeneracy curve maintaining the cell at a set target firing rate (see Fig (1C)). As the statistics of the neuron’s membrane potential change, the combinations of *β* and *h* required to preserve the target firing rate change too, repositioning the degeneracy curve in parameter space.

To demonstrate how intrinsic plasticity shapes the excitability profile of neurons (Fig (1)), simulations were performed in which individual neurons were exposed to different inputs (resulting from both connectivity and external stimuli). From these simulations, three categories (classed by ad hoc thresholds of impinging firing rate to each neuron) of excitability emerged as a result of neuron exposure to persistent high-, mid- or low-amplitude activity (Fig (1D)). For those neurons receiving high-amplitude activity, compared to baseline, the intrinsic plasticity-driven excitability changes in *h* (Fig (1E)) and *β* (Fig (1F)), yielded hypoexcitability. In contrast, low-amplitude activity did the opposite and yielded hyperexcitability. Despite displaying the same target firing rate, such changes in excitability result in an important difference in the neuron’s response to forthcoming stimuli, as both the rheobase (i.e., *h*) and slope (i.e., *β*) of the resulting excitability curve had changed significantly. These results echo the experimentally observed homeostatic responses of neurons to low and high activity [25, 26, 29], supporting the qualitative behavior of this model of intrinsic plasticity.

### B. Intrinsic plasticity supports changes in cell-to-cell heterogeneity and network dynamics

Our central hypothesis is that, within an active network, the tuning of cellular excitability through intrinsic plasticity will modulate population diversity. To investigate this potential relationship between diversity and activity, we placed our network of neurons into three distinct activity level regimes by applying a constant input equally to all neurons (i.e., +*/* −*I*_*o*_) reflective of three different physiological/experimental states: 1) a baseline state, where the network receives no external stimulation, and thus the activity of the population is entirely dictated by the recurrent connectivity of the network; 2) a high activity state similar to what is observed in the upstate of the cortex [83, 84] or in a region receiving feed-forward drive [85]; and 3) a low activity state similar to what is found in slice culture [86], or deafferented cortex [87] where the population receives a scarcity of excitatory drive.

The top row of Fig (2A) shows that persistent high and low activity states cause shifts in excitability compared to baseline, Fig (1), that one would expect experimentally [25, 26]. In the network, these changes are reflected in the shape of the mean network response function, ⟨*f* (*u, β*_*eff*_, *h*_*eff*_) ⟩ _*N*_. This mean network response function was computed by assuming that intrinsic plasticity operates over long time scales compared to ongoing membrane potential fluctuations, rendering the excitability profile of individual neurons essentially constant on shorter time scales. The mean response function is characterized by an effective parameterization (*β*_*eff*_ and *h*_*eff*_) of the network excitability, where the effective rheobase, *h*_*eff*_ is the midpoint of the population-average excitability curve (e.g. ⟨*f* (*u*_*j*_, *β*_*j*_, *h*_*j*_) ⟩ _*N*_ = 1*/*2) and the effective slope, *β*_*eff*_, can be determined from the derivative of this curve (see Methods). The value of these parameters reflects the heterogeneity in excitability found in the network and was determined numerically. We found that, in the high activity state, excitability was suppressed homeostatically (Fig (2Ai)) whereas in the low activity state neurons became more excitable (Fig (2Aiii)) [26] before returning to baseline after persistent high and low activity stops (bottom row of Fig (2Ai-iii)).

The parameterizations of the excitability, expectedly, change in opposing ways to adapt to the activity changes experienced with exposure to persistent high and low activity regimes (Fig (2B)). In line with Fig (1A-C), the neurons exposed to high activity adopt more hypo excitable curves, characterized by lower *β* and high *h* values, and the neurons exposed to low activity adopt more hyper-excitable curves, with higher *β* and lower *h* values (Fig (2B)). Concurrent with these excitability changes, the population firing rate also rose or dropped in response to high and low drive, respectively, but normalized over time back towards their homeostatic target via intrinsic plasticity (see Methods; Fig (2C,D)). Likewise, when the induced activity was turned off, the networks experienced opposing deviations in their firing rate before being driven back toward their homeostatic target again. The amount of time required for a given neuron in the network to regain the target firing rate is dependent on the excitability it possessed at the onset of high- or low-activity (Fig (2C)).

Beyond these predictable network-level excitability changes, surprisingly, these shifts in excitability were accompanied by a remarkable amount of malleability in heterogeneity (Fig (2Ai-iii)). Heterogeneity was quantified by computing the population-averaged Hellinger distance, ⟨*H*^2^⟩ _*N*_, (a similarity metric see Methods) to track the variability in excitability profiles among cells. Using this metric, the network demonstrated a decline in heterogeneity for both activity states. However, in the high-activity state, this initial loss recovers to an overall increase in heterogeneity as the activity persists (Fig (2E)). The initial nominal drop in heterogeneity in the high-activity state occurs as the sudden increase in activity induces intrinsic plasticity, and all the neurons start moving toward more hypoexcitable parameterizations. However, as the high activity persists, the interneuron variance in activity (e.g. from differing connectivity degree) brings the recovery and eventual overtake of the baseline heterogeneity. Moreover, the cessation of the high-activity state induces further heterogeneity. This differs from the low activity state, which experiences a profound decline in diversity, where the interneuron variability in activity becomes small owing to the suppressing nature of the applied input (see Methods). Interestingly, this lost heterogeneity is readily recovered when the low activity state stops.

In the low-activity state modeled here, and paralleled by our recent human slice culture experiments (ref), a substantial reduction in diversity is accompanied by shifts in mean excitability. These features are similarly reproduced in our simulations (Fig (2 Aiii)). While a simplified representation, our model effectively captures the adaptive regulation of diversity and its relationship to excitability through intrinsic plasticity mechanisms observed in biological neural networks, thus highlighting how diversity is maintained by activity, declining without it.

Recent experimental evidence, corroborated by numerical and analytical work, has associated reduced heterogeneity in neuron excitability with pathological states [13], and found it broadly destabilizing for brain activity [48, 49]. The results presented in Fig (2) supports these findings, showing that stimulation can be used to manipulate the excitability (i.e., Fig (2A)) and level of heterogeneity (i.e., Fig (2E)) of a population of neurons. It remains to explore the consequences of these heterogeneity changes on the network’s dynamics. To do this, we exposed networks endowed with the excitability curves obtained in the high-, low- and baseline activity states (Fig 2A, top row) to a slow, time-varying stimulus (i.e., *η*(*t*)) in the form of an Ornstein-Uhlenbeck process selected to interrogate the system’s dynamics (see Methods, Section II G 1). To avoid confounds, and because it occurred on a much slower time scale, intrinsic plasticity was temporarily turned off for this analysis.

The network response to the applied stimulus is shown in Fig (3A). In the baseline case, the neurons’ responses varied smoothly, mirroring the stimulation applied. Variability in spiking was observed amongst neurons, as a direct consequence of high excitability heterogeneity (Fig (2A)). However, both other regimes had stereotyped network responses: In the high-activity state, the network remained mostly silent and unresponsive, revealing both hypo-excitability and reduced heterogeneity. In the low activity state, the network dynamics were characterized by sharp, network-wide (i.e., identical between cells) fluctuations alternating between states of full quiescence and states of saturation. These are the signatures of the hyperexcitability and decline in heterogeneity depicted in Fig (2A). Such transitions emerged as the direct consequence of the increased non-linearity of the network mean response function. This homogenization-induced nonlinearity was especially important when the network was in the hyperexcitable regime, wherein bursts of intense activity resulted from small perturbations, suggestive of multistability. To further interrogate this result, we performed linear stability analysis on the network (see Methods and Fig (Supp_*F*_ *ig*)).

**FIG. 3.**
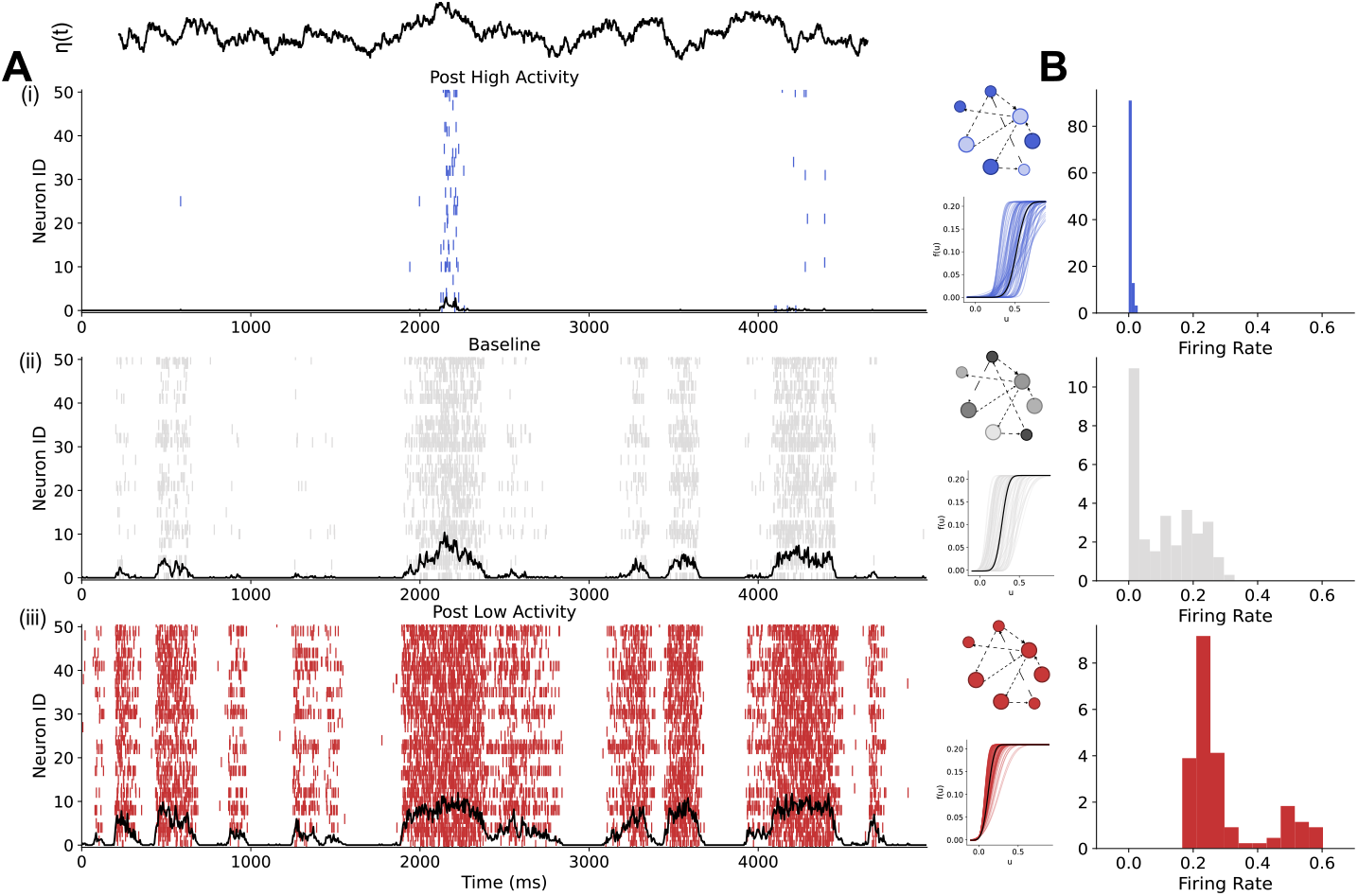
High- and low- activity induced excitability alterations yield aberrant network dynamics. Using the results from the chronic stimulation regimes of Fig (2A), we assess the effects of excitability on the stability of the network. With the model’s excitability curves frozen (*t*_*Fig2A*_ = 19900 ms) each of the three resultant networks was re-simulated for Δ*t* = 1000 ms with a new slow stimulus, *η*(*t*) (see Methods). **A)** Raster of a subset of the neuron responses to the new slow stimulus *η* (see Eq (37)). **B)** Distribution of the firing rates of the *N* = 100 neurons simulated in each network in (A).

Further in line with the spiking behaviour exhibited by the different networks, the high activity hypoexcitable neurons had the lowest firing rates, and the mean of the low-activity hyperexcitable neurons was highest (Fig (3)Bi-iii)). These firing rates are reflective of the excitability, showing the high (red) through low (blue) firing rates that result when such hyperexcitable or hypoexcitable populations receive input different from the persistent activity.

### C. Network heterogeneity reflects variability in cellular milieus

Our results indicate that excitability heterogeneity is an activity-dependent, malleable property of our network model. This malleability arises directly from the intrinsic plasticity equations through an implicit dependence on the membrane potential of individual cells. The impetuous for any alteration in the heterogeneity is thus supported by changes to its membrane potential statistics (e.g. ⟨*u*_*i*_⟩_*T*_ and 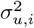) that define the neuron’s *milieu* here defined as the environment or local conditions that impact a neuron’s response [88] resulting from the combination of passive properties (e.g., membrane time constant, leaking term), synaptic inputs, and external stimuli. Thus, as has been hypothesized for cells in organ tissue [89, 90], we postulate that heterogeneity is a direct consequence of variability in cellular milieu wherein the neurons adapt to their local conditions. In our model, this is then manifested through intrinsic plasticity. To challenge this claim, we first considered factors that influence the firing rate, *r*_*i*_, and connectivity weights, *w*_*ij*_, of the neurons due to their direct impact on membrane potential mean (Eq (12)) and variance (Eq (13)).

To characterize how unique each neuron is from the population, we calculated its Hellinger distance 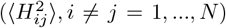 which represents the difference of neuron *i*’s excitability profile from those of the *N −* 1 other neurons in the network. This was performed on the network in its baseline regime (one where no stimulation is present; *I*_*o*_ = 0). This resultant “uniqueness” metric was plotted with respect to the first two moments of the membrane potential of individual cells, that is the temporal mean, ⟨*u*_*i*_⟩_*T*_ (Fig (4A)), and variance, 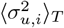 (Fig (4B)).

**FIG. 4.**
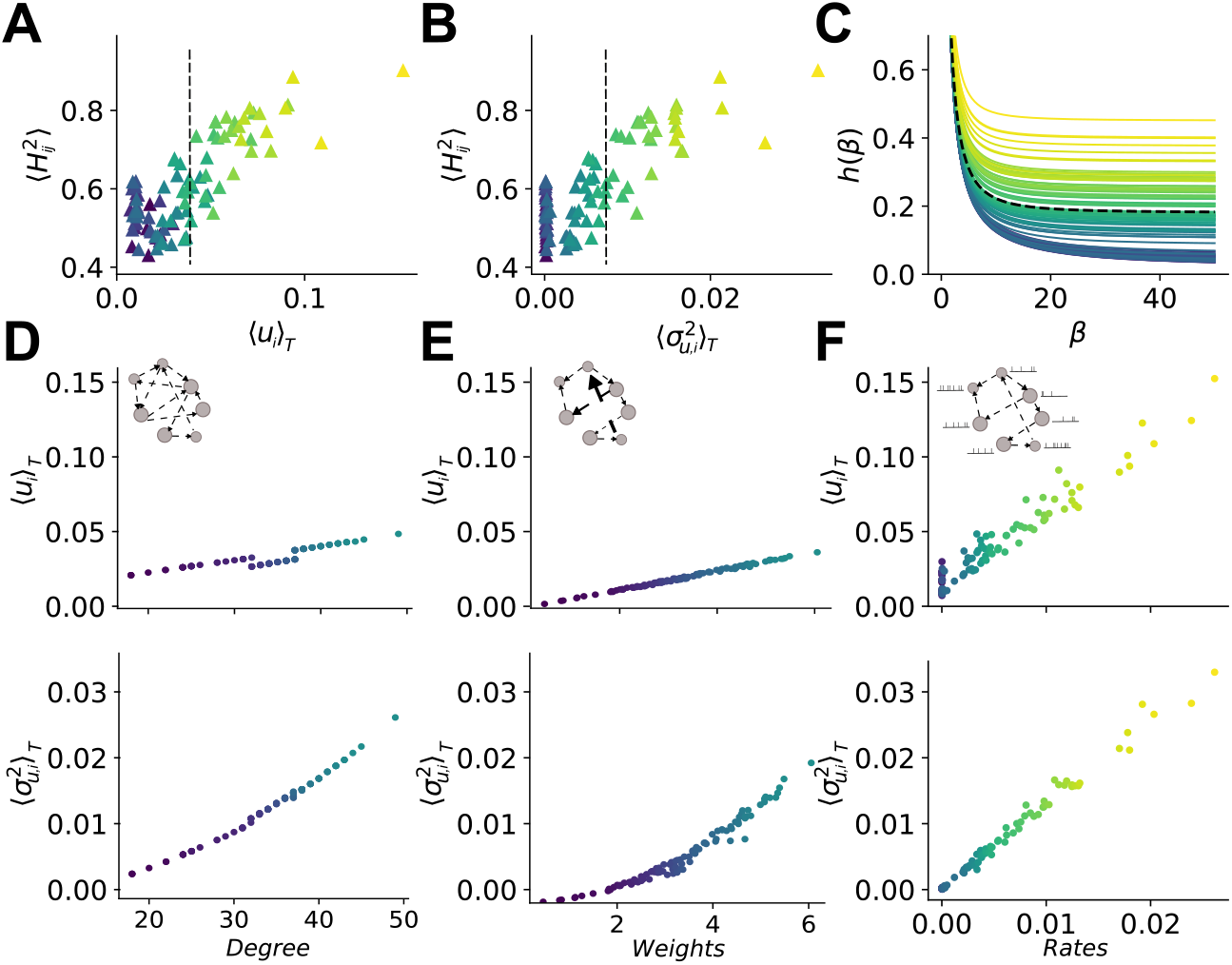
Network properties affecting membrane potential statistics regulate population diversity. The average of the pairwise Hellinger distance for each neuron, 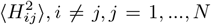 as a function of the membrane potential **A)** mean, ⟨*u*_*i*_⟩ _*T*_, and **B)** variance, 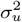, of each neuron, *i* = 1, …, *N*. The population membrane potential mean, ⟨*u*⟩ _*N*_ and variance, 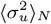 are shown as dashed vertical lines in each plot. **C)** The degeneracy curves of the neurons are plotted in (A) and (B). The dashed degeneracy curve is found using the population mean and variance (⟨*u*⟩ _*N*_ and 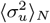). Each degeneracy curve’s colour matches the neuron’s point in plots (A) and (B). The mean (top) and variance (bottom) of the membrane potential of neurons for varying **D)** in-degree, **E)** sum of the realized weight, *W*, and **F)** rate of stimulation

We found that the neurons with the most unique excitability profiles were those with the most unique milieus - characterized by statistics far from the population average (dashed lines in Figs (4A,B)). This was also reflected in their degeneracy curves (Fig (4C)), where those of high ⟨*u*_*i*_⟩_*T*_ and 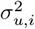 were positioned far from the population average curve (dashed line). The separation of these degeneracy curves from those around the population average will, via intrinsic plasticity, lead to very different parameterizations of their excitability (Fig (1C)) and hence contribute to amplify heterogeneity in the network. These results indicate that factors which increase the variability of membrane potential statistics between cells - i.e., diversify the milieus - promote heterogeneity. To understand the underpinning factors that influence the mean and variance of the membrane potential, which define the milieu, we considered three factors that have major influence on single cell milieus as per Eq. (12) and (13): the network topology via input degree (Fig (4D)), the connectivity weight (Fig (4E)), and the presynaptic firing rate, *r*_*s*_, (Fig (4F)). The latter acts as a proxy for a case where the impinging firing rates could be directly manipulated. In line with their theoretical relationship, the mean of the membrane potential were found to scale linearly with the degree of connectivity, synaptic weights and presynaptic firing rate, and scaled non-linearly with the variance of the synaptic weights and connectivity.

Having characterized a relationship between milieu features and the excitability of individual neurons, we sought to elucidate the contributions of these features to network heterogeneity. To do this, we performed a series of simulations where the variances of each trait were systematically increased for different distribution means. At the end of the simulation, the network’s effective excitability (Fig (5), top row) and heterogeneity (Fig (5), bottom row) were calculated as the population-average Hellinger distance (see Methods).

**FIG. 5.**
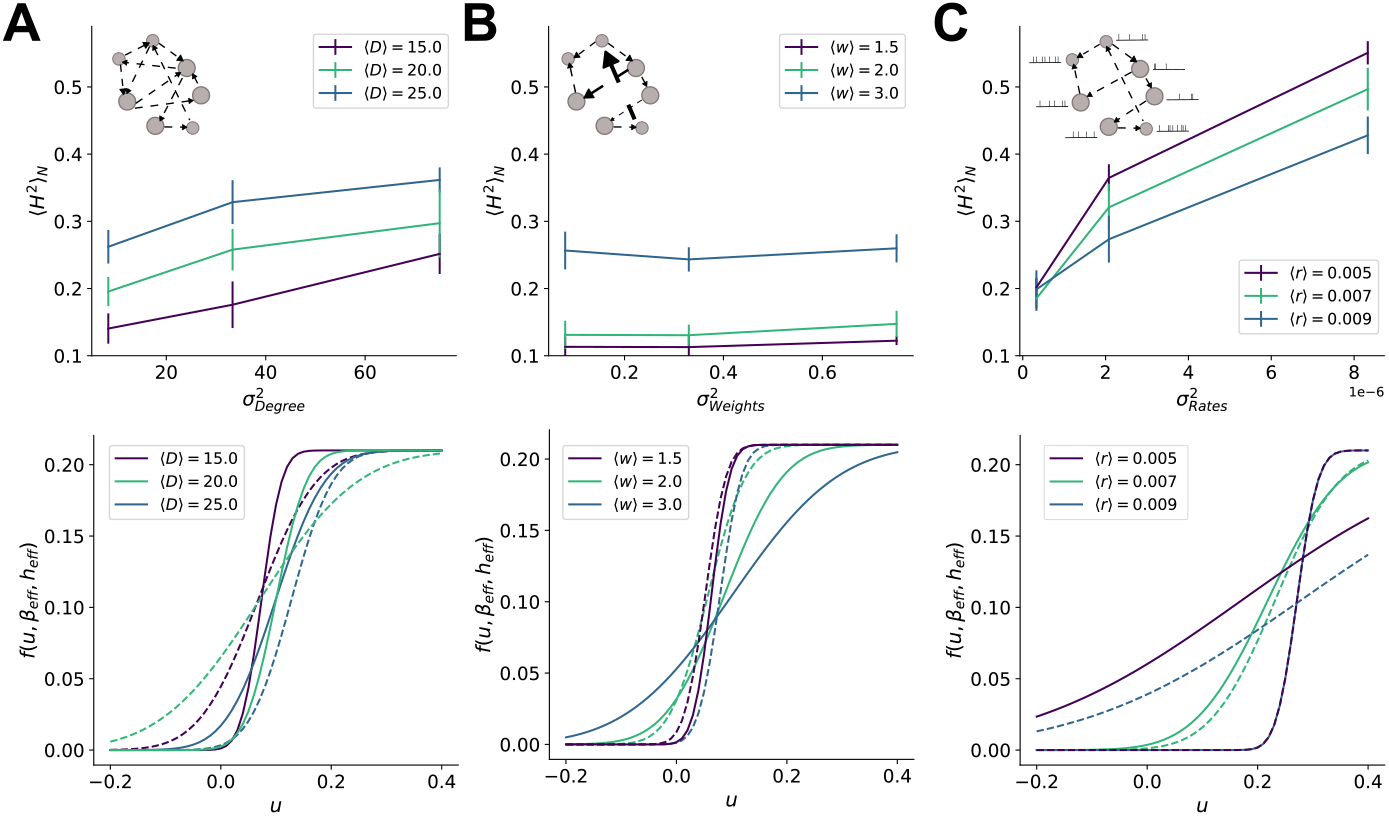
Increased variance of degree distributions and stimulation rate but not weights yields elevated hetero-geneity. For networks of increasing variance in **A)** the degree distribution, 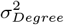, **B)** the weight distribution, 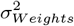, and **C)** the rates of Poisson spike trains that act as surrogates for presynatpic firing rate (see Methods),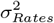, population-average heterogeneity, ⟨*H*^2^⟩ _*N*_ (top), and the effective excitability profiles (bottom) are plotted for three different means. For the effective excitability profiles, the dashed lines correspond to the lowest variance cases and the solid lines to the highest variance cases. Error bars are the standard deviation across ten independent simulations at each variance level. When not otherwise stated, the network for all subsequent simulations has a degree distribution with mean 15 connections and fixed weight, *w*_*ij*_ = *w*_*o*_, for all connections. The network is simulated for *T* = 20000 ms, during which intrinsic plasticity allows the evolution of each neuron’s excitability in response to the neuron’s milieus.

Beginning with the degree of connectivity (Fig (5A)), we found that regardless of the mean degree, increasing the variance of the degree distribution increased the heterogeneity of the network. The observed increase in heterogeneity from high ⟨*Degree*⟩ and 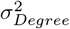 was in agreement with our analytical predictions, where ⟨*u*_*i*_⟩_*T*_ (Eq (12)) and 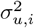), and hence the milieu, were related to degree as it controls the number of local synaptic connections. In contrast, Fig (5B) showed that changes in effective excitability scaled mostly with the mean synaptic weight, ⟨*w*⟩, but did not vary significantly with the distribution’s variance, 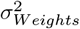. Not unexpectedly increasing the mean of the degree distribution, ⟨*Degree*⟩, produced a more hypoexcitable effective excitability. Increasing the mean of the weight distribution similarly increased the network heterogeneity. However, increasing the variance of the weight distribution had no such effect (Fig (5B)). Finally, we investigated the influence of the presynpatic firing rate on excitability and heterogeneity. As with degree, the heterogeneity of the network increased with higher variance in the distribution of presynpatic firing rates (Fig (5C)). Taken together, these results suggest that degree and presynaptic firing rate are the most influential on the neurons’ milieus, and hence the heterogeneity of the network.

### D. Application: external stimulation as a candidate for stabilizing changes in heterogeneity

Collectively, the previous results indicated that: 1) intrinsic plasticity makes heterogeneity malleable; 2) variability in milieu features promotes increased diversity in the network; and 3) such increased diversity is activity-dependent. We therefore sought to explore the malleability of heterogeneity in the context of neuromodulation, where exogenous stimulation may be a potential therapeutic intervention. Specifically, we wished to verify 1) whether external stimulation may be used to modulate the heterogeneity of our network model and 2) if such induced changes in heterogeneity might persist in time (i.e., whether networks can retain such stimulation-induced heterogeneity). To this end, we repeated the simulations of networks with different variances in the degree distribution (Fig (5A)), but included a fixed input, *I*_*o*_, injection for an epoch of duration Δ*t* = 10000 ms in the middle of the stimulation. From these simulations, we compare the heterogeneity in the network in three epochs: before (pre; Δ*t* = 15000 ms), during (stimulation; Δ*t* = 10000 ms), and after (post; Δ*t* = 10000 ms). The pre-stimulation duration is longer to allow the network excitabilities to reach a steady state from their randomized initialization. The heterogeneity of each of these epochs is measured by calculating ⟨*H*^2^⟩ _*N*_ for the network at a time point near the end of each epoch (*t* = 14000, 24500 and 34000 ms, respectively). This process was repeated for injections of input with increasing amplitude.

As can be seen in Fig (6A), heterogeneity increases with stimulation, *I*_*o*_, of sufficiently high amplitudes. This was true across a range of degree distribution variances. However, increasing variance of the degree itself had only limited effect, contributing only when stimulation was a moderate amplitude (Fig (6B)). This suggests, in agreement with Fig (5A,C), that more heterogeneity is recruited by the manipulation of presynaptic input than the variance of degree. Interestingly, the induced heterogeneity was found to persist after stimulation offset when the amplitude of stimulation was sufficiently large (Fig (6B)). In line with Fig (2), this suggests that the network, perhaps by recruiting the pre-existing heterogeneity from the distributed degree of connectivity, has enough variability to sustain the heterogeneity induced by stimulation.

**FIG. 6.**
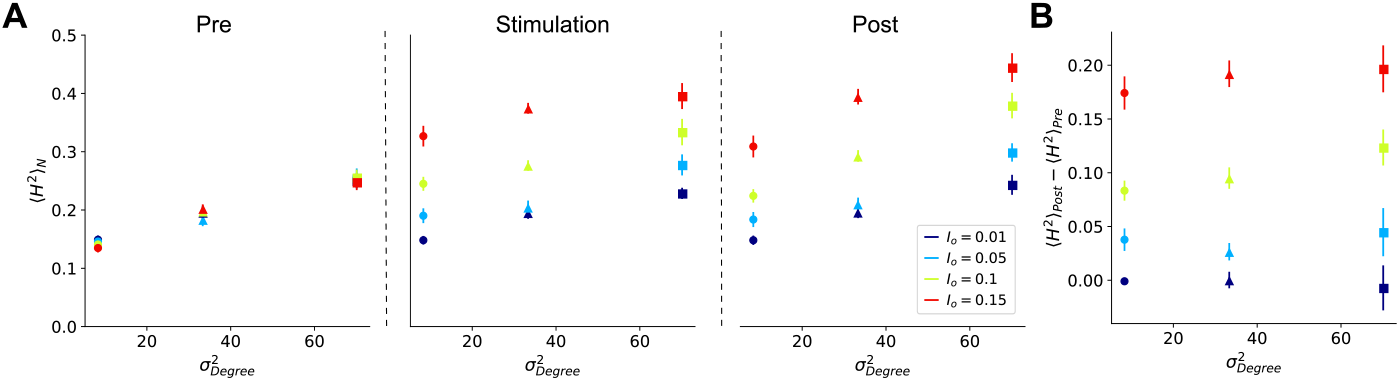
High amplitude stimulation induces heterogeneity across multiple variances in degree distributions. **A)** Three networks with increasing variances in their degree distributions and a mean ⟨*Degree*⟩ = 15, were simulated for *T* = 35000 ms. Each network received fixed-input injections, *I*_*o*_, at four different amplitudes for Δ*t* = 10000 ms (*t* = 15000 ms to *t* = 25000 ms). This was repeated 10 times, with the connectivity randomly drawn at the start of each trial. The heterogeneity of each simulation was calculated at three different time points: *t* = 14000 ms (pre-stimulation), *t* = 24500 ms (during), and *t* = 34000 ms (post-stimulation), as the population-average Hellinger distance, ⟨*H*^2^⟩ _*N*_. Error bars are SEM. **B)** The difference in the population-average Hellinger distance pre- and post-stimulation obtained in (A).

## IV. DISCUSSION

Our computational modelling and mathematical analyses reveal that neuronal activity “milieus” interacting with degenerate homeostatic intrinsic plasticity rules can actively modulate neuronal diversity. Herein, our work animates the conceptually static image of transcriptomic and electrophysiological diversity [Eq. (12), Eq. (15), Eq. (16), Eq. (51), Eq. (52), Eq. (54), Eq. (91), Eq. (92)], towards understanding within cell-type diversity as a homeodynamic process. In this light, rather than a fixed monolithic property of neuronal populations, within-cell-type diversity takes on an adaptive property of networks dependent on activity statistics, connectivity, and intrinsic plasticity rules. This has particularly salient implications, as recent genetics research has found that neurons receiving different inputs (e.g. baroreceptor versus catecholaminergic) form unique clusters in reduced dimensional anaylsis [89]. That is, differential inputs to individual cells drive population variability at the phenotypic level.

Crucially, in our network model, this occurred in neurons endowed with a degenerate intrinsic plasticity mechanism, wherein qualitative similarity of their dynamics (e.g. firing rate) is achieved with diverse excitability curves. Such degeneracy has been observed throughout the nervous system, including at the level of ion channels [67, 68, 93], which directly relates to the plasticity rule implemented here. Including degeneracy in our framework distinguishes the approach from existing adaptation models [Eq. (60)–Eq. (65), Eq. (76), Eq. (77)], where neurons exposed to the same input statistics reach identical parameterizations. The intrinsic plasticity in the model serves to modulate the level of diversity in response to the milieus the neurons experience. However, in actual neurons, changes in excitability manifest as biophysical alterations such as upregulation of ion channels [18, 53, 76, 88]. By including the ability to capture these differences in our model, we obtained a much more biophysically plausible model [68, 79, 93, 94], supporting the candidacy of intrinsic plasticity as a mechanism by which diversity is actively maintained, and which can be lost under pathological conditions.

Our finding that the low-activity state gives rise to seizure-like dynamics through diversity decline is particularly compelling, as it supports the emerging view that network ‘silence’ may serve as a convergent pathway to seizure generation across diverse neurological and neurodegenerative disorders [84]. Although such silence is thought to destabilize network dynamics through homeostatic increases in excitability alone [25, 26, 84], our previous work suggests the associated diversity decline is equally culpable in allowing pathological dynamics to emerge [13, 48, 49].

Aside from its utility for pathological explorations, our model’s relative simplicity allowed us, with analytical motivation, to interrogate what network properties support heterogeneity. Specifically, we consider those features that effect the membrane potential statistics, ⟨*u*_*i*_⟩_*T*_ and 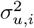, (Eq. (12) and (13) and Fig (4)) that comprise each neuron’s milieu. In our analysis, we found each neuron’s milieu was defined by the presynaptic firing rate, connectivity, and weights. Increasing the mean of any of the three properties and increasing the variance of the degree or presynaptic firing rate directly increased network heterogeneity (see Fig (5)). The network weights having less influence on the ability of the network to induce heterogeneity implies that the local milieu differences between neurons acquired by varying the weights are small compared to those created by topology (e.g. degree). This is of particular interest as it has been reported in recent years that the probability of connectivity within and between different cortical layers and regions is variable [16, 95]. This may then serve a purpose in bolstering the heterogeneity within the brain. That is, in line with transcriptomic work [89] and theory in non-neuronal cells [90], increasing the variability in input to individual neurons shifts the network into near-subclasses reflective of the milieus the neurons experience (e.g. separation of neurons with low degree and hence membrane potential mean and variance from those with a high degree). This separation of milieus begins to cause increased diversity in the excitability analogous to what is observed under different activity levels as was simulated (see Fig (1D-F)) and aligns with experimental manipulation of presynaptic firing rate [29].

In this context, it is well established that the number of synaptic connections a neuron receives — its degree — is log-normally distributed, reflecting the heavy-tailed structure of cortical and hippocampal networks[96]. Interestingly, transcriptomic levels of many ion channel genes also exhibit log-normal distributions. Our work here provides a parsimonious explanation for this convergence via intrinsic plasticity [97]. Neurons with high synaptic degree tend to fire more frequently and with higher burst probability, engaging calcium-dependent signalling cascades and transcriptional regulators such as CREB and Fos, which in turn modulate ion channel gene expression [98]. Because both synaptic input and transcriptional output are shaped by multiplicative, synergistic processes—rather than linear summation—their resulting distributions naturally follow log-normal statistics [96]. This link provides a mechanistic bridge from network topology to cellular identity, suggesting that intrinsic plasticity may serve to reinforce or even amplify the functional stratification imposed by the log-normal architecture of connectivity.

The malleability of heterogeneity observed in our model shares a remarkable convergence with our previously reported experimental papers [13]. This is particularly true in the low activity state (see Fig (2Aiii)), which paralleled our recent human slice culture experiments [47] where a substantial reduction in diversity is accompanied by shifts in mean excitability. In our computational approach, the low activity state is induced by applying a negative fixed input (see Methods). Similar to a slice experiment, wherein the neurons are deafferented, this pushes the neurons in our network to a ∼ homogenous, near-zero mean that will move the target parameters of the neurons into the hyperexcitable regime (see Fig (1C) and Methods). That is, the milieus of the neurons become aggressively similar. This differs from the high activity state, where applying a positive fixed input yields an initial transient decline in heterogeneity (see Fig (2E)). However, ultimately, the baseline variance in the milieus created by the network connectivity is preserved (unlike the large quenching experienced in the low activity regime) and pushes the neurons to new, greater levels of heterogeneity across the hypoexcitable regime. Compared to physiological regimes, which are constrained by factors such as energy expenditure, our model has no upper bound in parameteric space.

From a functional perspective, our current work positions the work of Tripathy et al. [11] in a “dynamic” light, suggesting that heterogeneity can be tuned by the statistics of afferent input. Their results show that population coding is optimized not by maximal heterogeneity, but by an intermediate level tailored to the structure of the stimulus. This suggests that intrinsic plasticity mechanisms could adapt the diversity of a neuronal population to match its functional demands. For example, a population exposed to highly periodic, low-dimensional input — such as a rhythmic oscillation — may converge toward similar intrinsic properties, effectively homogenizing the population to encode redundant temporal/spatial features efficiently. In contrast, exposure to more asynchronous, high-entropy input, rich in temporal or spectral complexity, could diversify the population, broadening its receptive field space to enhance coding capacity. An excellent example of this is the visual system that is pluri-potent in the activity it can generate in reponse to stimuli of varying spatial statistics. We have provided computational evidence of how, within the same network, visual input of low spatial variance generates gamma oscillations, whereas stimuli with high spatial variance generate broadband power changes indicative of asynchronous (high-entropy) population activity [99]. Thus, stimulus-driven patterns of activity may shape population heterogeneity through intrinsic plasticity, enabling neural circuits to flexibly allocate variability in service of efficient coding.

### A. Limitations

Due to the mobile nature of the degeneracy curves in response to increases in ⟨*u*⟩ and 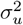, and the equations which regulate the intrinsic plasticity (Eq (3) and (4)), the network is sensitive to its initialization conditions. If the initialization of *β* and *h* starts a sizable portion of the neurons on or near the multistable region (see Fig (7C) shaded region) or in the hyperexcitable regime it may result in initially high activity that will prompt a period of hypoexcitability in the network by causing a large upward migration of the degeneracy curves. Further, the lack of inhibitory activity in the model predisposes the network to bursts of high activity if a sufficient portion of the neurons occupies the hyperexcitable parameter space at the same time. Such states result in intermittent fluctuations in the rate as any sudden burst of high activity induces the network’s plasticity to adapt toward hypoexcitability.

**FIG. 7.**
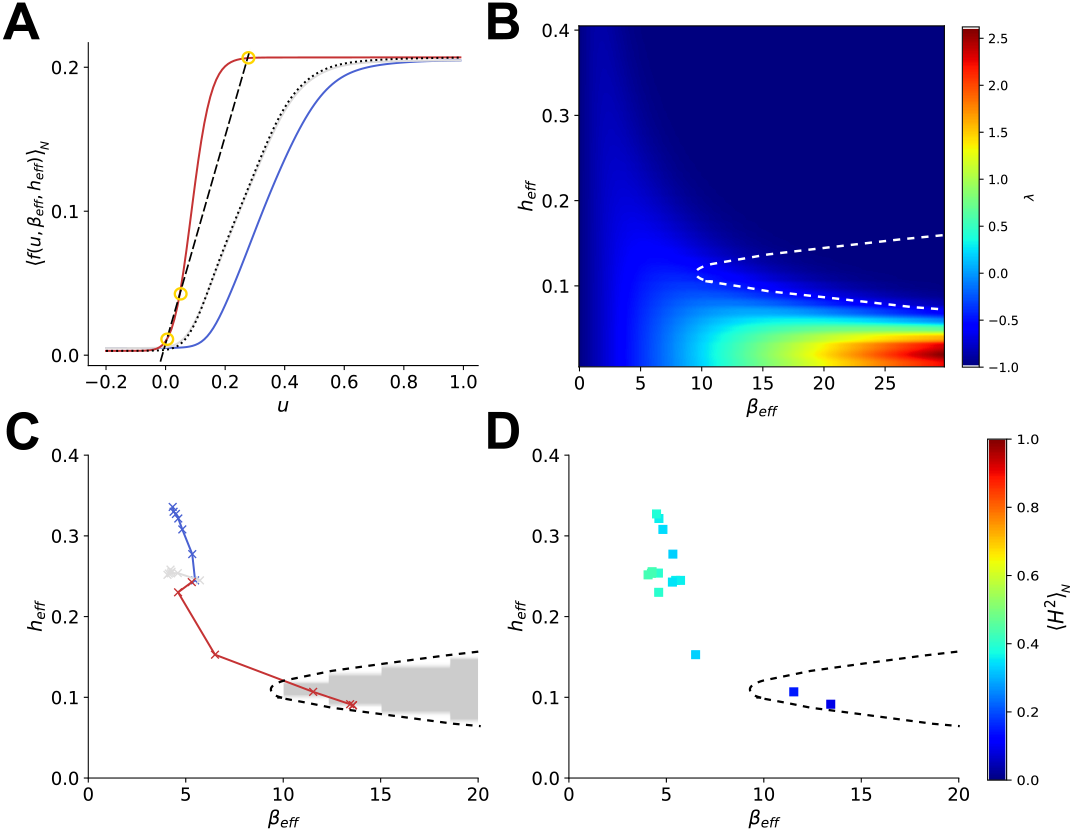
Linearization and stability analysis. **A)** The population average excitability profile, ⟨*f* (*u, β, h*) ⟩ _*N*_, for the network under chronic over- and under-stimulation and at baseline as simulated in Fig (2A). The dotted line is ⟨*f* (*u, β, h*) ⟩ _*N*_ prestimulation. Yellow circles indicate fixed points of the red excitability curve. The dashed line in these plots represents the graphical intercept indicating the fixed-point location(s) on each excitability curve. **B)** The effective parameterizations of ⟨*f* (*u, β, h*) ⟩ _*N*_ over the time course of the simulation. The shaded region is the graphically determined multistable regime. The dashed line around this region is replicated on (C) and (D) for visualization. **C)** The eigenvalue of the network, *λ*, as calculated from the mean field predictions of the model (see Methods; Eq (31)) for combinations of *β*_*eff*_ and *h*_*eff*_ when the weights of the network are *w*_*o*_ = 0.21 and the target firing rate is *r*^*o*^ = 0.05. **D)** Relationship of the heterogeneity level in the network, measured as ⟨*H*^2^⟩ (see Eq (36)), to the effective excitability.

To quantify the amount of heterogeneity in the networks here, we use the Hellinger distance as a metric (see Methods). We use this in order to have a metric by which we can directly, analytically relate the parameters *β* and *h* to the heterogeneity, in this case via their relationship to the mean and variance of the derivative of the excitability curve. Importantly, though both parameters contribute to the heterogeneity measured by this metric, there is a disproportionate weighting to heterogeneity in *β* compared to *h*. This is due to the scale difference in the parameters and the variance of *β* being accounted for by a squared term (see Eq (36)). As a result of this, if the heterogeneity is increased, for example, before and after stimulation predominantly in *h*, it may appear small in ⟨*H*^2^⟩ for such cases. This can be verified, visually, instead by looking at the standard deviations of the excitability curves when overlaid.

## V. CONCLUSION

We have demonstrated that a degenerate intrinsic plasticity rule is a viable mechanism for dynamically regulating heterogeneity in an excitatory network. Specifically, we showed that intrinsic plasticity can sustain, increase, or decrease diversity. Such “dynamic diversity” is dependent on the milieus experienced by the neurons in the network as a function of the input statistics to each neuron, which are manipulated by external stimuli, and amount of diversity itself. These results thus form the framework for future investigation into how the statistics that arise in more complex networks may influence the heterogeneity and hence functional capacity [11] and resilience of neuronal networks [13].

## VI. AUTHOR CONTRIBUTIONS

**Conceptualization:** Daniel Trotter, Jeremie Lefebvre, Taufik Valiante

**Funding acquisition:** Jeremie Lefebvre

**Formal Analysis:** Daniel Trotter, Jeremie Lefebvre

**Investigation:** Daniel Trotter

**Methodology:** Daniel Trotter, Jeremie Lefebvre

**Supervision:** Jeremie Lefebvre

**Validation:** Daniel Trotter, Jeremie Lefebvre

**Visualization:** Daniel Trotter, Jeremie Lefebvre

**Writing - original draft:** Daniel Trotter

**Writing - review and editing:** Daniel Trotter, Tau-fik Valiante, Jeremie Lefebvre

## VII. ACKNOWLEDGEMENTS

We thank the New Frontiers in Research Fund (Grant NFRFE-2023-00354 to JL) and the National Research Council of Canada (Grant RGPIN-2017-06662 to JL) for support of this research.

## VIII. DATA & CODE AVAILABILITY

The code used for this study can be found at https://github.com/djtrotter/intrinsic_plast.

## Appendix A

