## Supplementary figures and images for "Intrinsic plasticity underlies malleability of neural network heterogeneity"

### Supplemental Figure 1

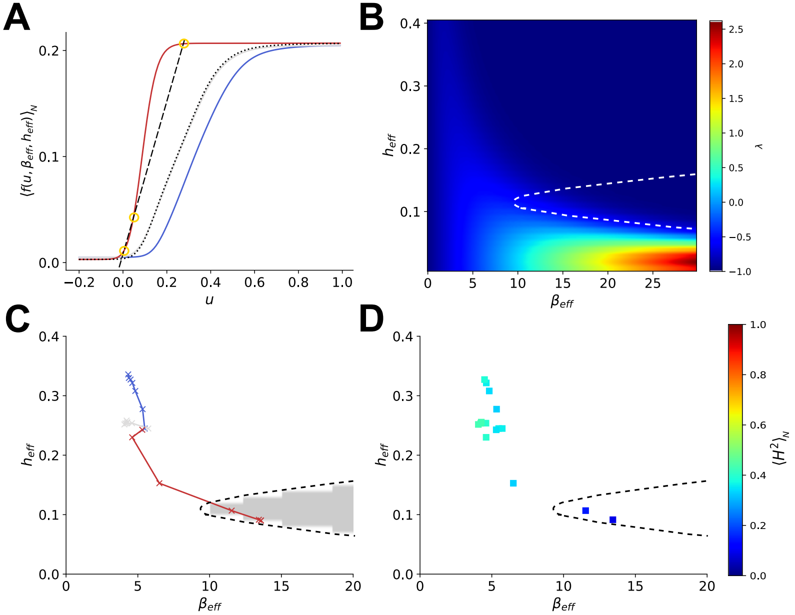
